# *ARL15* promotes inflammatory fibroblast activation and disease severity in rheumatoid arthritis: integrated transcriptomic and collagen-induced arthritis model analyses

**DOI:** 10.64898/2026.06.26.733622

**Authors:** Sujit Kashyap, Anuj Kumar Pandey, Manisha Saini, Kumari Vijaya, Sriram Kunnoth, Puneet Mahajan, Suman Kundu, Uma Kumar, B.K. Thelma

**Affiliations:** Department of Genetics, University of Delhi South Campus, New Delhi 110021; Department of Biochemistry, University of Delhi South Campus, New Delhi 110021; Department of Computer Science, Birla Institute of Technology, Ranchi 835215; Department of Applied Mechanics, Indian Institute of Technology, Delhi 110016; Department of Rheumatology, All India institute of Medical Sciences New Delhi 110016

## Abstract

**Background:** ADP-ribosylation factor-like protein 15 (*ARL15*) is a rheumatoid arthritis (RA) susceptibility gene identified through GWAS. Previous studies suggested a role for ARL15 in synovial fibroblast (SF) pathogenicity, but its contribution to inflammatory arthritis remains unclear. We investigated the inflammatory role of ARL15 and its therapeutic potential in RA.

**Methods:** *ARL15* was overexpressed in MH7A cells followed by bulk RNA sequencing and pathway enrichment analyses. Therapeutic relevance was evaluated in collagen-induced arthritis (CIA) mouse model using anti-ARL15 monoclonal antibodies, *ARL15*-targeting siRNA, or isoquinoline. Arthritis scores, histopathology, micro-CT and serum cytokines were assessed. Publicly available single-cell RNA sequencing (scRNA-seq) datasets were analyzed to determine *ARL15* expression in RASF subsets.

**Results:** *ARL15* overexpression induced a pro-inflammatory transcriptional program characterized by upregulation of *IL1A, IL1B, IL6, IL8, CXCL1, CXCL10*, and *CCL20*. Gene set enrichment analysis revealed activation of *IL6–JAK–STAT, TNF*, interferon-response, and KRAS signaling pathways, with suppression of oxidative phosphorylation, lipid metabolism, and mTORC1 signaling. In CIA mice, ARL15 inhibition significantly reduced arthritis severity, inflammatory infiltrates, and joint destruction while preserving cartilage and bone integrity. Serum TNF-α, IL-6, and IL-1β levels were markedly decreased following ARL15 blockade. Combination monoclonal antibody treatment demonstrated the greatest therapeutic benefit. scRNA-seq analysis showed broad *ARL15* expression across RA fibroblast populations, with enrichment in inflammatory lining and SF subsets.

**Conclusions:** ARL15 is a pro-inflammatory regulator of SF activation and arthritis progression. Integrated transcriptomic, single-cell, and *in vivo* analyses identify ARL15 as a therapeutic target for RA and support further translational development of ARL15 based therapies.

## 1. Introduction

Rheumatoid arthritis (RA) like several other common complex traits is a clinically and pathogenetically heterogeneous disease. With its etiology being complex and elusive, this autoimmune condition continues to present therapeutic challenges. Nonsteroidal anti-inflammatory drugs (NSAIDs) are the first line of treatment for RA patients in early disease stage. However, swelling of the feet, heartburn, stomach ulcers, possible risk of blood clot and heart attack are some of the side effects observed with NSAIDs(1). Disease-modifying antirheumatic drugs (DMARDs) are the next favored group for treatment. Of these, Methotrexate (MTX) is the DMARD of choice but only ∼ 60% of RA patients respond to this drug. Use of MTX as well as other DMARDs like sulfasalazine, hydroxycholoroquine are also associated with leucopenia, bleeding gums and increased risk of infections warranting alternate treatment regimens(2), and the focus shifted to biologicals. Cytokines have been shown to play a major role in the pathogenesis of this autoimmune disorder with TNFα emerging important in inflammation induced joint damage in RA. Levels of major inflammatory cytokines including TNFα, IL6, IL1β and IL17 have been measured to be higher in RA patients as compared to healthy individuals(3). Thus, pro-inflammatory cytokines are attractive targets paving the way for the next generation drugs for RA. This group of drugs commonly referred to as biologicals is very specific for the targets, requiring fewer doses to be administered as compared to DMARDs. Introduction of etanercept and other biologicals targeting inflammatory cytokines and immune cell subsets have indeed improved the outcome and quality of life of patients with RA(4). However, due to their high costs and consequent non affordability combined with major side effects such as injection site reactions, redness and swelling, infusion reactions, difficulty in breathing, nausea, vomiting, rapid or weak pulse and high risk of serious infections like TB, the search for better drugs continues(5). Identification of new druggable targets is thus an activity of tremendous importance attracting academicians and pharma companies.

On the other hand, the series of candidate gene-based as well as the hypothesis free genome-wide association (GWA) testing of RA across ethnic populations have yielded a plethora of risk conferring genes. Most of these genes are functionally relevant and are a good resource for furthering our understanding of disease biology and possible development of new therapeutic molecules(6). ADP ribosylation like protein 15 (*ARL15*) is one such novel GWAS hit identified in the first ever GWAS in RA among the genetically distinct north Indian population(7). *ARL15* is a small GTPase and belongs to ADP ribosylation family (ARF). This family of proteins regulates vesicle trafficking and modulates membrane lipid composition(8). RA is closely associated with CVD risk factors. Several GWASs have found *ARL15* to be associated with metabolic traits such as HDL cholesterol concentration, insulin resistance, coronary heart disease, fasting insulin and triglyceride concentrations(9). *ARL15* has also been shown to be associated with age of onset for alcohol dependence(10) which in turn is correlated with risk of RA(11).

ARL15 has been shown to enhance AKT phosphorylation in HEK cells and thus affect PI3K-AKT pathway(9). Of note, PI3Ks (Phosphoinositide-3-kinases) are lipid sensing kinases that control cellular growth, proliferation, survival and apoptosis. *In vitro* studies have confirmed that the decreased invasiveness of fibroblasts is mediated by reduced phosphorylation of AKT and extracellular signal-regulated kinase(12). Interestingly, we have demonstrated reduced rheumatoid arthritis synovial fibroblast (RASF) invasion and migration along with downregulation of *IL6* in *ARL15* knockdown in RASF(13). Furthermore, transcriptome analysis following *ARL15* knockdown in RASF captured lowered expression of extracellular matrix stabilizer gene *COMP* which is linked to severe RA and an enhanced expression of adiponectin and IFN response genes namely *IFI6* and *USP18*. In addition, upregulation of genes implicated in disease modulation/treatment response; and downregulation of genes (*CTGF, CD248*, and *PTX)* suggestive of *ARL15* involvement *in* inflammation and RA-associated cardiovascular risk were noted. GO analysis of the findings showed PI3K-AKT pathway, a known druggable target for RA to be significantly affected. Conversely, *ARL15* KD in MH7A cells showed upregulation of cytokines (*IL1A, IL8, CXCLs*) and downregulation of inflammatory regulators (*DOCK2, TLR4, TGFB2*)(14). These observations are suggestive of *ARL15* being a potential therapeutic target and encourage additional investigations including animal studies.

Experimental animal models of arthritis provide important tools for the development and evaluation of new therapeutic approaches. Collagen-induced arthritis (CIA) is one such experimental model of arthritis induced in susceptible strains of mice by immunization with heterologous collagen type II, a joint-specific protein, in complete Freund’s adjuvant (15). As in the case of RA, CIA is primarily an autoimmune disease of the joints with increased angiogenesis, inflammatory cell infiltration, synovial hyperplasia, and bone erosion. Murine CIA has been frequently used to investigate mechanisms relevant to RA as well as new antiarthritic treatments(16). For example, SecinH3 is reported to inhibit all cytohesins which activate ARF(17). It is interesting to note that CIA mouse model of RA treated with SecinH3, notably improved the arthritis index by blocking MYD88–ARNO–ARF6 pathway, which was comparable with the Enbrel treatment of CIA mice (18). Taken together the suggestive modifier role of *ARL15* in RA, in this study we explored i) the likely mechanism of *ARL15* action through an *in vitro* overexpression strategy; and ii) its druggability using CIA mouse model and inhibitors.

## 2. Methodology

Further *in vitro* functional characterization by overexpression of *ARL15* and establishing its potential druggability to alleviate RA symptoms in CIA mouse model was explored.

### 2.1 Overexpression studies - MH7A cell line

#### Cloning, MH7A cell line transfection, transcriptome sequencing and Bioinformatics

ARL15 expression plasmid (Origene, USA) was amplified in E. coli DH5α, verified by restriction enzyme digestion and Sanger sequencing, and transfected into MH7A synovial fibroblast-like cells using Lipofectamine 2000. Cells were harvested 48 hours post-transfection for RNA isolation. Transcriptome sequencing was performed on ribodepleted RNA libraries using the Illumina NovaSeq 6000 platform (150 bp paired-end reads). Raw sequencing data underwent quality control and alignment to the human reference genome (hg19) using HISAT2. Gene-level read counts were generated using FeatureCounts and normalized with DESeq2. Differentially expressed genes were identified using a significance threshold of p ≤ 0.05 and |log2 fold change| ≥ 1, followed by downstream pathway enrichment analyses. Detailed experimental procedures are provided in the **Supplementary Methods**.

### 2.2 Animal studies

All guidelines provided by the Committee for Purpose of Control and Supervision of Experiments on Animals (CPCSEA), India; with study design and procedures approved by the Institutional Animal Ethics Committee (IAEC) of Eurofins Advinus Limited (Approval # 11_EAL_108_Renewal_Collagen arthritis_(CIA) in DBA1J mice, May 2021), where the study was conducted, were followed.

DBA/1J male mice 6-8 weeks old obtained from Jackson Laboratory, USA were clinically examined for good health and suitability for generation of the CIA model for the study. Each mouse was identified by an animal accession number, cage card and permanent tail marking during treatment phase. Mice were acclimatized for five days in experimental room before start of the experimental procedures. During this period, animals were observed at least once daily for any abnormalities. Of the total of 155 mice, n=143 were immunized; and n=12 maintained as naïve. Of the 143 animals, 96 developed the desired phenotype and were included in the study. Remaining 47 animals were humanely killed. A total of nine treatment groups with five test compounds and appropriate controls were used for this study (Supplementary Table 1). The standard protocol was used for immunization and scoring for the disease symptoms/severity (see supplementary material, Figure 1A).

**Figure 1.**
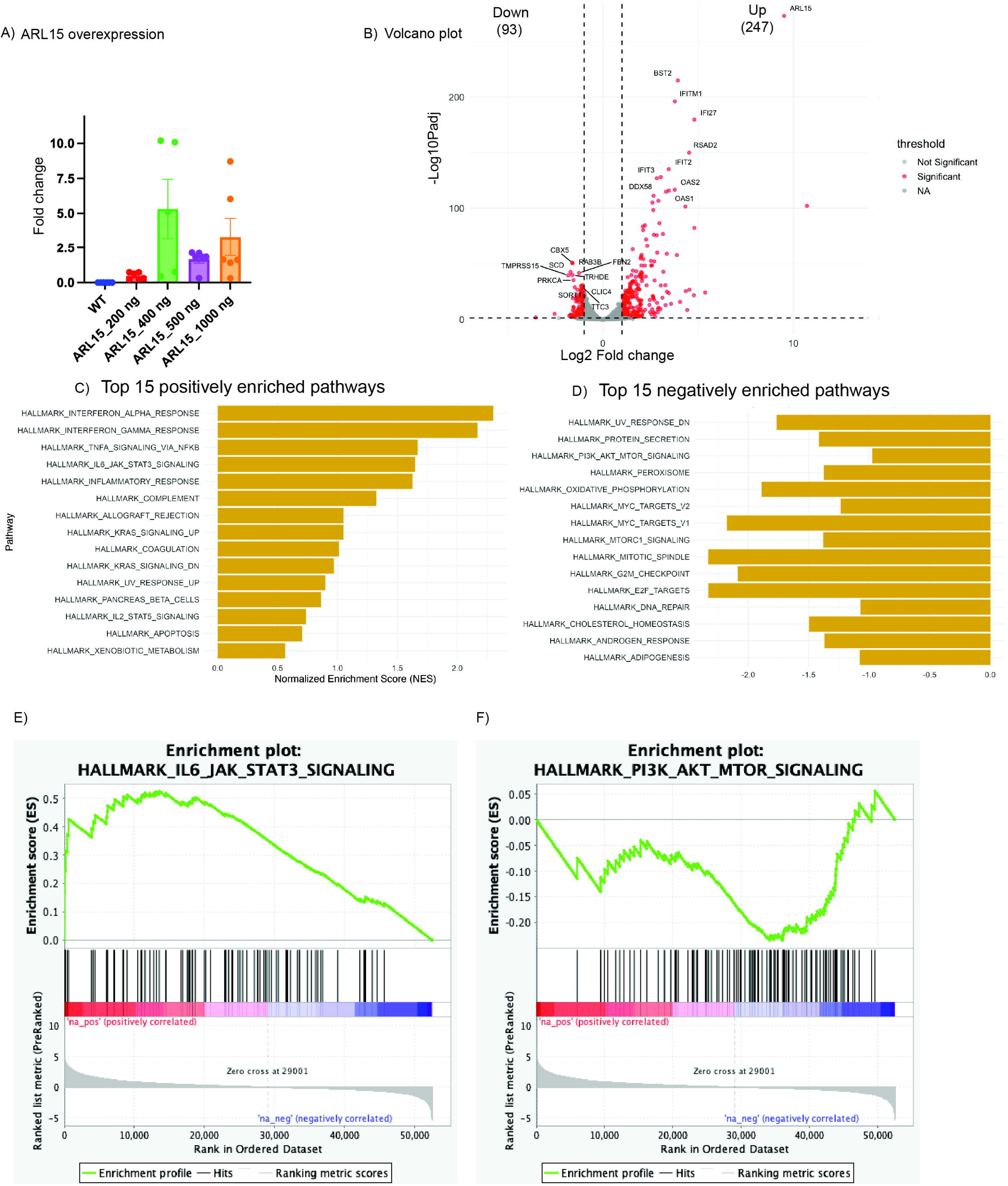
Over expression in MH7A. Over expression of ARL15 in MH7AA) qPCR plot showing expression of ARL15 at various plasmid concentration. B) Volcano plot showing the genes differentially regulated upon ARL15 overexpression. C) & D) shows pathways enriched by genes up and down regulated upon ARL15 overexpression in MH7A cells. E) & F) shows enrichment plot of IL6 jak-stat signaling and Pl3-AKT signaling from genes up and downregulated respectively by ARL15 over expression.

#### Inflammatory marker estimations

Serum biomarkers namely IL-6, TNFα, IL-1β, IL-17, collagen immunoglobulins IgG1 & IgG2a were quantitated using ELISA. Percentage change between each treatment group were calculated and relevant graphs and tables plotted (see supplementary Material).

#### Histopathology

Histopathology of synovial joints was documented for all the treatment groups. The severity of arthritis was classified as Minimal, Mild, Moderate & Severe based on H&E staining according to the following criteria: Minimal: synovitis, without cartilage loss and bone erosions. Mild: synovitis, cartilage loss, and bone erosions limited to discrete foci Moderate: synovitis and erosions present but joint integrity maintained. Severe: synovitis, extensive erosions and loss of joint integrity.

The severity of proteoglycan loss was classified as minimal, mild, moderate, or severe based on Safranin-O staining according to the following criteria: Minimal: Minimal loss; Mild: Mild loss; Moderate: Moderate loss; Severe: Severe loss.

#### Micro CT scanning of mouse knee joints and paws

Micro CT scanning was done in the laboratory of the collaborator Professor Puneet Mahajan, IIT, New Delhi. Of a total of ten mice bones (3 mice each from CIA, Isotype control and ARL15 mAb1 groups and a single healthy control mouse), was performed at 70kVp, 114µA, (irrespective of the group), by exposing the bones to radiation for 300ms, with formalin as the medium. Only hind limbs of mice were used for this analysis. Scanning at 10µm resolution was performed using 0.1 aluminium filter to reduce the beam hardening effect.

#### Image Processing

Once the scan was completed, the in-built reconstruction algorithm was used to develop the cross sectional images. These images were processed using image processing technique to develop the 3D images of mice bones. The range of global threshold, 654.2-1239.7 mg HA/ccm, was used to develop the 3D geometry of the bones.

### 2.3 Single-cell RNA sequencing dataset acquisition and preprocessing

Publicly available single-cell RNA sequencing (scRNA-seq) data from rheumatoid arthritis (RA) synovial tissue were obtained from ImmPort under study accession SDY2213 (19). Data from five RA patients were included in the present analysis. The analysis workflow and preprocessing strategy were adapted from the published pipeline available in the RA Fibroblast Multiome Analysis GitHub repository. Additional customized scripts were implemented in R using the Seurat framework for downstream visualization and expression analysis.

Quality-controlled fibroblast-like synoviocytes (FLS) were integrated and visualized using Uniform Manifold Approximation and Projection (UMAP). Fibroblast subsets were annotated according to the previously defined fine-grained synovial fibroblast classifications, including Inflamed lining, Inflamed sublining, Intermediate inflamed/resting lining, Intermediate lining/sublining, Resting lining, Resting sublining.

#### Gene expression analysis

Expression of *ARL15* was assessed across all fibroblast subsets using normalized scRNA-seq expression values. UMAP feature plots were generated to visualize the spatial distribution of *ARL15*-expressing cells across synovial fibroblast populations. To compare expression among fibroblast subsets, violin plots and dot plots were generated. Dot plots summarized both the proportion of expressing cells and the average expression level within each subgroup. To further evaluate the inflammatory context associated with *ARL15* expression, feature plots for the inflammatory mediators *IL6* and *TNF* were also generated and compared with the *ARL15* expression pattern.

#### Statistical analysis

All data obtained as detailed above were collated and quantities are presented as the mean and analyzed using the GraphPad Prism Version 5 software program (GraphPad Prism, San Diego, CA, USA). Group comparisons were performed using t-test and One-way ANOVA nonparametric tests, and statistical significance was recorded with P<0.05.

## 3. Results

### 3.1 Over expression analysis

Overexpression of *ARL15* in MH7A was confirmed using qPCR (Figure 1A). Differential expression analysis between control and overexpressed MH7A cells showed 247 genes upregulated (Supplementary table 2) and 93 genes downregulated (Supplementary table 3) in MH7A cells with *ARL15* significantly overexpressed (Figure 1B). *ARL15* overexpression strongly induced pro-inflammatory cytokines and chemokines. Notably, *IL1A, IL1B, IL6*, and *IL8* were significantly upregulated, indicating activation of classical inflammatory pathways (Figure 1C). A wide range of chemokines involved in immune cell recruitment were also increased, including *CXCL1, CXCL2, CXCL3, CXCL6, CXCL10, CXCL11, CCL5, CCL20*, and *CCL26*. In contrast to the strong induction of immune-related genes, *ARL15* overexpression resulted in significant downregulation of genes involved in lipid metabolism and biosynthesis. These included *ACACA, FADS2, SCD, HMGCS1*, and *DHCR24*, which are key regulators of fatty acid and cholesterol synthesis (Figure 1D). This pattern indicates metabolic reprogramming consistent with an activated inflammatory state, in which anabolic lipid pathways are often suppressed. GSEA with upregulated genes showed hallmark enrichment of IL6 JAK Stat signaling (Figure 1E), TNF signaling, interferon response and KRAS signaling. GSEA with downregulated genes showed hallmarks enrichment of Mtorc1 signaling (Figure 1F), cholesterol homeostasis and oxidative phosphorylation.

### 3.2 Visual evaluation of arthritis in CIA mouse model

Animal groups immunized with collagen showed symptoms of disease progression such as inflammation in joints within 12-16 days of primary immunization. Visual examination showed that there was significant loss of body weight in CIA mice compared with control mice (Supplementary figure 2A); severity of disease as measured by arthritis score was also found to be increased in CIA mice as compared to control mice before the start of treatment (Supplementary Table 4 & 5, Figure 2B). After administration of all the treatment doses within 14 days, the redness and swelling of CIA and Isotype controls remained unchanged but mAb, siRNA and Isoquinoline treated group showed decrease in disease severity.

**Figure 2.**
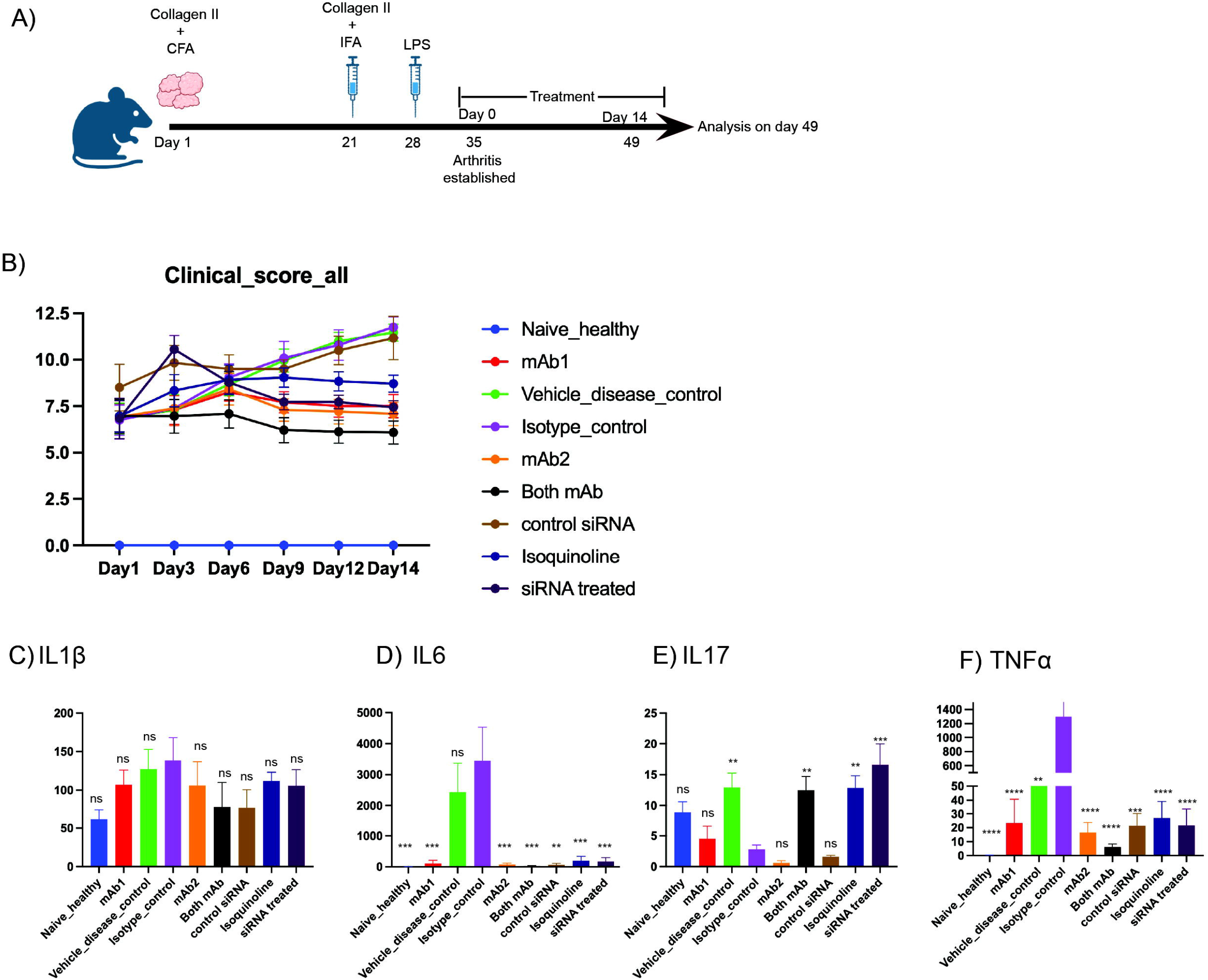
ARL15 inhibition in CIA mice model. A) Schematic diagram of treatment of mice in the CIA model B) Clinical score of all the mice used in this study over two weeks C) to F) Measurement of serum cytokine levels. From left to right IL1, IL6, IL17 and TNFa. Data are presented as mean ± SEM. Statistical analyses were performed using one-way ANOVA followed by Dunnett’s multiple-comparison test against the lsotype control group. *P<0.05, **P<0.01, ***P<0.001, ****P<0.0001

**Clinical Score**: The Vehicle (Disease control) group showed progressive arthritis, with mean total clinical scores of 6.83 on Day 35, rising to 11.46 on Day 49. Arthritis scores for anti-ARL15 treated groups decreased significantly in 2 weeks (Figure 2B, Supplementary Table 5). **Paw Thickness**: The mean hind paw thickness in the Naïve control was 1.750 mm. The Vehicle group displayed increased paw thickness (2.631 mm). The group treated with AntiARL15 antibody mAb 1 showed a lower mean paw thickness (1.897 mm) compared to the Vehicle and Isotype Control groups (Supplementary Table 6).

### 3.3 Micro CT radiographic evaluation

CIA mice with no treatment or treatment with isotype control showed higher bone degradation evidenced by the black marks in the scanned images. ARL15 mAb treated mice showed much lesser damage as compared to the CIA and isotype control groups in the paw and knee joints. Healthy mice scanned images were clean with no damage marks (Figure 4).

### 3.4 ARL15 blockade attenuates synovial inflammation and joint destruction in CIA mice

Histopathological analysis of ankle joint sections from collagen-induced arthritis (CIA) mice demonstrated marked inflammatory and destructive changes compared with healthy controls. Healthy mice exhibited normal joint architecture with intact cartilage and bone structures and no evidence of inflammatory cell infiltration (Figure 3A). In contrast, vehicle-treated CIA mice showed severe synovial hyperplasia, dense inflammatory infiltrates, extensive pannus formation, and pronounced bone erosion, confirming successful induction of arthritis (Figure 3B). Similarly, mice treated with isotype control antibody or control siRNA displayed persistent inflammatory pathology characterized by synovial expansion, invasive pannus tissue, and substantial cartilage and bone destruction, indicating no therapeutic benefit from control treatments (Figure 3C, G).

**Figure 3.**
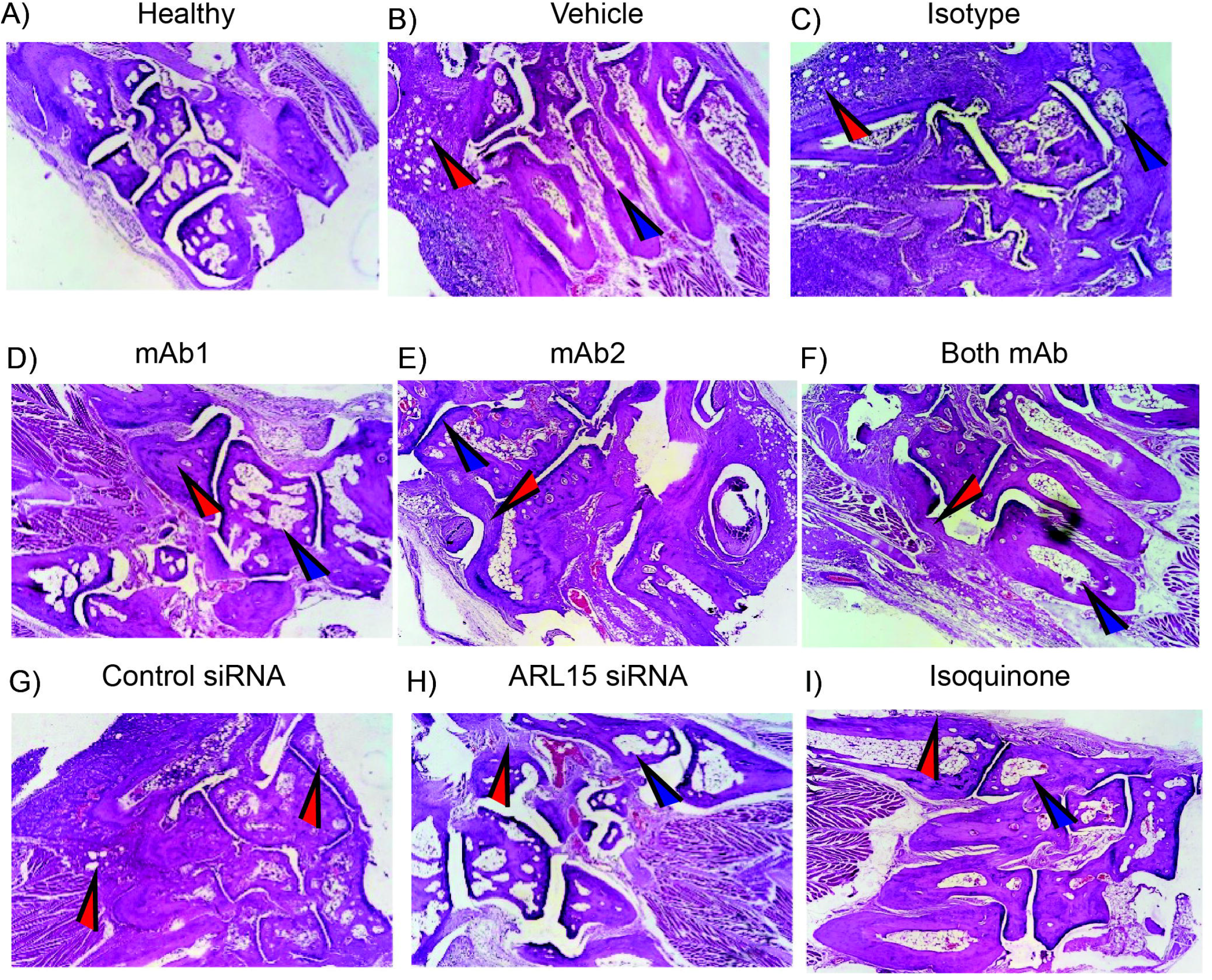
H&E staining with healthy mice. H & E staining analysis of bone joints at 4X magnification. From left to right, bone joints of mice from healthy, untreated (PBS), lsotype treated and monoclonal antibody treated. Blue arrow indicating bone erosion and red arrow indicating synovial inflammation with pannus formation.

**Figure 4.**
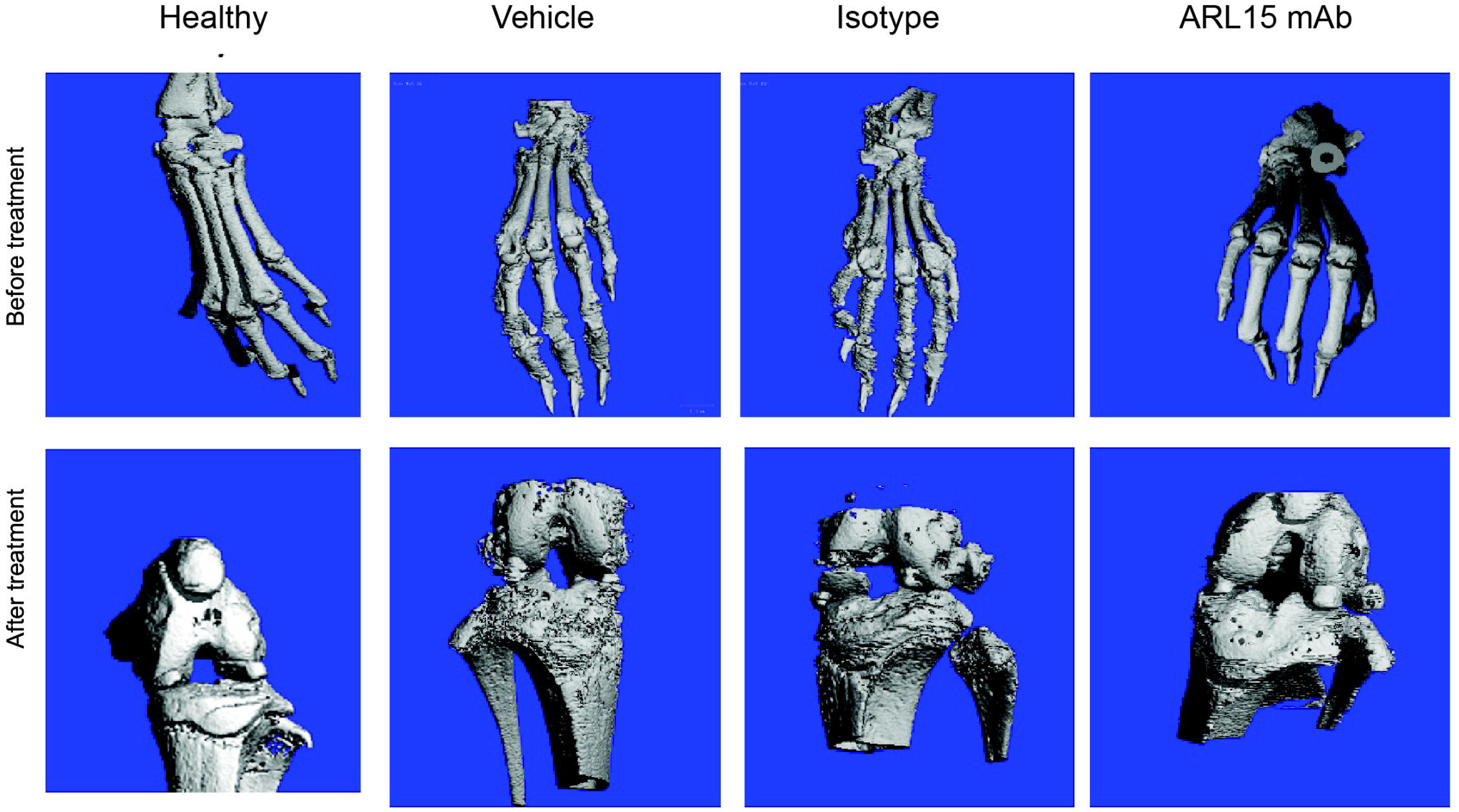
Micro-CT imgaing. Micro-CT analysis of bone joints before and after treatment. From left to right, bone joints of mice from healthy, untreated (PBS), lsotype treated and monoclonal antibody treated.

Treatment with monoclonal antibodies (mAb1 or mAb2) reduced the severity of arthritis-associated histopathological changes (Figure 3D, E). Both treatments decreased inflammatory cell infiltration and pannus formation while partially preserving joint integrity and bone structure. Notably, combined treatment with both monoclonal antibodies produced a greater protective effect, with markedly reduced synovial inflammation and minimal bone erosion compared with vehicle-treated CIA mice (Figure 3F). Silencing of *ARL15* using siRNA also significantly ameliorated joint pathology (Figure 3H). *ARL15* siRNA-treated mice exhibited reduced inflammatory infiltrates, decreased pannus formation, and preservation of cartilage and bone architecture relative to control siRNA-treated animals, suggesting a pathogenic role of *ARL15* in CIA progression. In addition, isoquinoline treatment attenuated joint destruction and inflammatory responses, as evidenced by reduced synovial inflammation, diminished pannus formation, and improved preservation of bone morphology (Figure 3I).

**Naïve Control (G1)**: Showed Normal Hind Paw pathology (NAD) and an H&E score of 0. **Vehicle Control (G2)**: Showed signs of Arthritis, Mild in H&E staining and Proteoglycans loss, Minimal in Safranin-O staining (Supplementary figure 2D), with a mean H&E score of 1.33. Isotype Control (G3): Showed Arthritis, Moderate in some animals (Animal 27, 28) and a mean H&E score of 2.17. AntiARL15 antibody mAb 1 (G4): Showed the best histopathological result, with 4 out of 6 animals showing NAD (No Abnormality Detected) and a mean H&E score of 0.17. This score was significantly lower than the Isotype Control. siRNA (G8) and Isoquinoline (G9): Both groups also showed improved scores (H&E Mean: 0.50) compared to the Isotype Control.

### 3.5 ARL15 inhibition suppresses systemic inflammatory cytokines in CIA mice

Serum cytokine analysis demonstrated marked reduction of IL-6, TNF-α upon anti-ARL15 treatments. Levels of IL-1β was also reduced, although not significantly but and IL-17 showed an increase in some treated groups in CIA mice compared with healthy controls. Treatment with anti-ARL15 monoclonal antibodies showed the most prominent effect on circulating inflammatory cytokines, with combination antibody therapy producing the greatest suppression of IL-6, TNF-α, and IL-1β. ARL15-targeting siRNA and isoquinoline treatment also reduced inflammatory cytokine levels compared with disease controls. Detailed cytokine measurements including mean SEM values are provided in supplementary material and supplementary table 7.

### 3.6 *ARL15* is expressed across RA synovial fibroblasts populations

Integrated scRNA-seq analysis of synovial fibroblasts from five RA patients identified distinct fibroblast subpopulations corresponding to inflamed and resting lining and sublining states (Figure 5A). UMAP visualization demonstrated clear segregation of fibroblast subsets, consistent with previously reported synovial fibroblast heterogeneity. Feature plot analysis showed that *ARL15* was broadly expressed throughout the fibroblast compartment, with detectable expression across both lining and sublining populations (Figure 5B). Notably, elevated expression was observed in inflamed fibroblast subsets, particularly within inflamed sublining and intermediate inflammatory populations. Dot plot analysis demonstrated that *ARL15* expression was present in a substantial proportion of cells across all fibroblast subtypes (Figure 5C). The inflamed lining and inflamed sublining populations exhibited both a higher fraction of *ARL15*-positive cells and increased mean expression levels relative to resting fibroblast populations. Violin plot analysis further confirmed widespread *ARL15* expression across fibroblast subsets, with comparatively higher median expression observed in inflammatory fibroblast states. In contrast, resting lining and resting sublining fibroblasts demonstrated relatively lower expression intensity (Figure 5D). Next, we attempted to find if *IL6* expression matched with the *ARL15* expression and we found them to be expressed in the inflamed sublining (Figure 5E).

**Figure 5.**
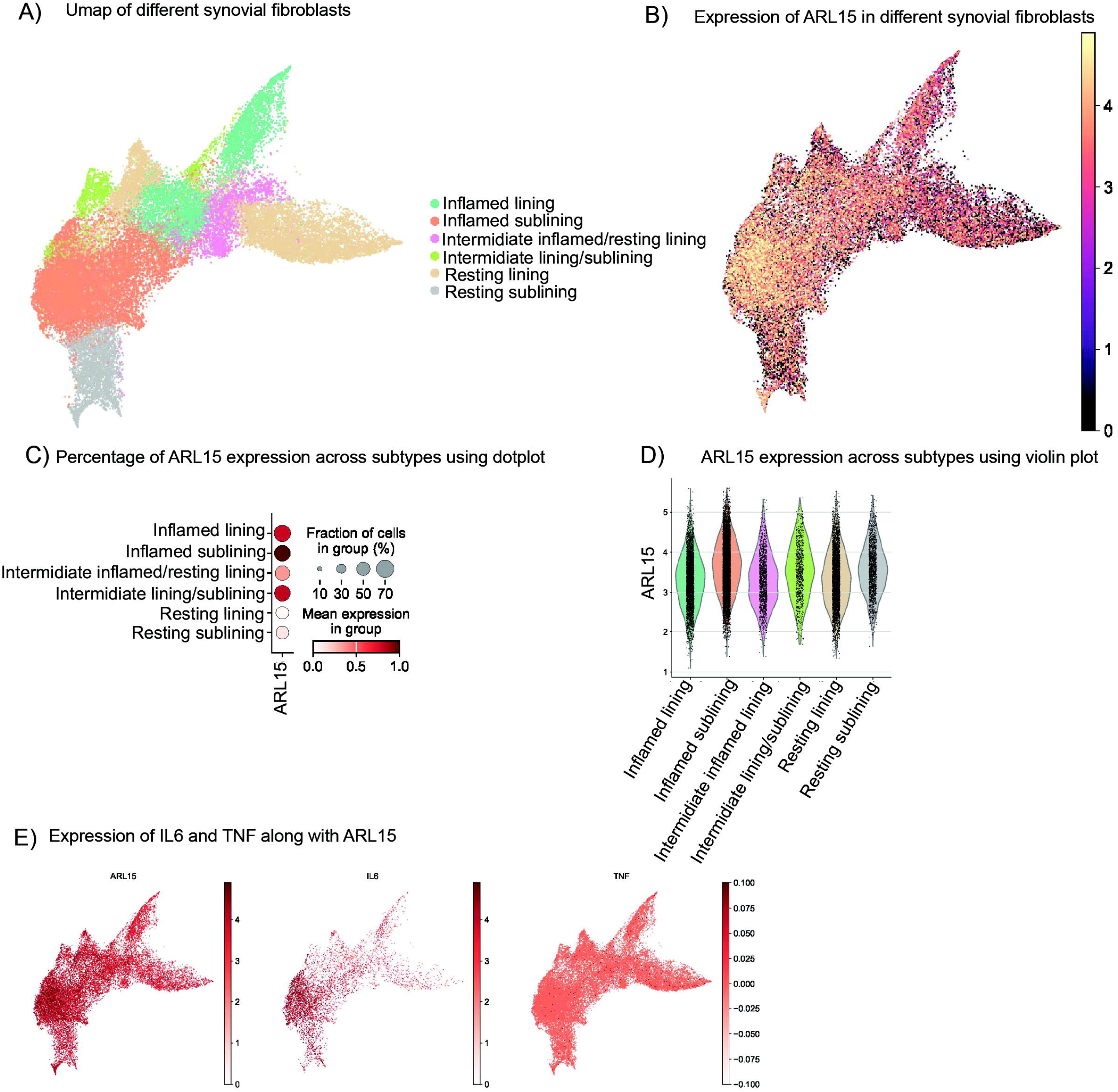
scRNAseq from synovial fibroblasts from RA patient synovium. scRNAseq from synovium of RA patients A) Umap showing all the subcluster beloning to various types of synocial fibrobla 8) ARL15 expression across all subtypes C) Dot plot showing the percentage of ARL15 expression per subtype Violin plot showing the percentage of ARL15 expression per subtype E) Umap showing the expression of inflamatory markers such as IL6 and TNF with along with ARL15.

## 4. Discussion

Preliminary characterization of ***ARL15***, a GWAS based susceptibility gene for RA, its likely modulatory role in disease, suggestive of its putative druggability have been documented in the work previously done in the laboratory (7, 13, 14, 20). To further these leads, three additional approaches namely i) overexpression of *ARL15* in MH7A cells; ii) scRNAseq analysis for *ARL15* expression using RA patient data; and iii) *in vivo* relevance of *ARL15* in CIA mice were explored in this study.

Overexpression of *ARL15* in MH7A synovial fibroblasts induced a robust inflammatory transcriptional program. Bulk RNA-seq followed by GSEA demonstrated strong positive enrichment of the inflammatory response, interferon-α, and interferon-γ hallmark pathways (NES > 1.5–2.0). These signatures encompassed classical interferon-stimulated genes and inflammatory mediators, indicating that elevated ARL15 levels can potentiate inflammatory signaling in fibroblast-like synoviocytes. Concomitantly, *ARL15* overexpression resulted in significant negative enrichment of cell-cycle hallmarks, including E2F targets, G2/M checkpoint, and mitotic spindle pathways (NES < –1.5) (Figure 1F). These findings suggest that ARL15 dampens proliferative programs in MH7A cells while enhancing inflammatory transcriptional states. Although MH7A is an immortalized, transformed fibroblast line, and thus may exhibit exaggerated interferon responsiveness to overexpression-driven cellular stress, the directional impact of ARL15 on inflammatory pathways is clear.

In primary (patient derived) RA synovial fibroblasts (RASFs), *ARL15* knockdown produced a distinct yet complementary phenotype, characterized by markedly reduced migration and invasion (13). These processes are central to pannus expansion and joint destruction, and are hallmarks of activated, tumor-like RASFs. The suppression of this invasive behavior upon ARL15 depletion indicates that endogenous ARL15 is required for maintaining the aggressive migratory phenotype of pathogenic fibroblasts. The divergence between proliferation-associated pathways in MH7A cells and motility-associated phenotypes in primary RASFs is consistent with the notion that MH7A and primary fibroblasts emphasize different effector modules of *ARL15* biology; importantly, both models converge on the concept that ARL15 enhances disease-relevant fibroblast activation.

The *in vivo* relevance of *ARL15* was confirmed using the CIA mouse model, where pharmacologic or genetic blockade of *ARL15* led to a notable reduction in arthritis scores, joint swelling, and inflammatory infiltrates (Supplementary Table 5, Figure 2B). These observations indicate that ARL15 is not merely a marker of fibroblast activation but is functionally required for efficient propagation of synovial inflammation and tissue damage *in vivo*. The attenuation of disease severity upon *ARL15* inhibition tested through multiple agents including antibodies, siRNA and small molecule (Figure 1B) supports a model in which ARL15-driven inflammatory and invasive fibroblast programs substantially contribute to RA pathogenesis. A major finding of the present study is that ARL15 inhibition markedly attenuated systemic inflammatory responses. The most striking effects were observed for IL-6 and TNF-α, both of which were reduced by more than 90% following anti-ARL15 antibody treatment. Of the inhibitors tested, a combination of two antibodies seemed to yield the best therapeutic effect., suggesting enhanced target blockade and supporting a direct role for ARL15 in regulating inflammatory signaling pathways. Both control and *ARL15*-targeting siRNA treatments were associated with reduced circulating IL-6 and TNF-α levels relative to disease controls. However, *ARL15* siRNA treatment produced greater improvements in histopathological and clinical disease parameters. In contrast to the substantial reductions observed in IL-6 and TNF-α, serum IL-1β levels were not significantly altered following ARL15-targeted interventions. IL-1β production is predominantly regulated through inflammasome activation in myeloid cells and macrophages, whereas our transcriptomic analyses identified ARL15 as a regulator of synovial fibroblast-associated inflammatory programs characterized by activation of IL6–JAK–STAT and TNF signaling pathways. The absence of significant changes in IL-1β therefore suggests that ARL15-mediated pathogenic effects are more closely linked to fibroblast-driven inflammatory networks than to inflammasome-dependent mechanisms (21, 22). This interpretation is further supported by the concordance between transcriptomic findings and the marked reductions in IL-6 and TNF-α observed following ARL15 inhibition in vivo. Unexpectedly, IL-17 levels were not reduced following ARL15 inhibition and showed a modest increase in some treatment groups despite significant improvements in clinical arthritis scores, histopathology, and bone preservation. IL-17 is a key pro-inflammatory cytokine implicated in synovial inflammation, osteoclastogenesis, and cartilage destruction in rheumatoid arthritis through its synergistic interactions with TNF-α and IL-1β (23, 24). However, previous studies have demonstrated that modulation of individual inflammatory pathways does not always result in parallel reductions in IL-17(25), suggesting the existence of compensatory immune mechanisms and pathway redundancy within the inflammatory network. The observed increase in IL-17 may therefore reflect compensatory activation of Th17-associated responses following suppression of dominant inflammatory mediators such as TNF-α and IL-6. Importantly, the marked therapeutic benefit observed following ARL15 blockade indicates that the pathogenic effects of ARL15 are primarily mediated through inflammatory pathways other than IL-17.

Taken together, the integration of these experimental approaches supports a coherent biological model in which ARL15 acts as a pro-pathogenic regulator that amplifies inflammatory transcriptional signaling and promotes the invasive behavior of synovial fibroblasts, thereby exacerbating joint inflammation and destruction. The inflammatory gene expression signatures observed in MH7A cells following *ARL15* overexpression are directionally consistent with the decreased inflammation observed in the CIA model upon ARL15 blockade (Figure 1; Supplementary Tables 2 and 3). Similarly, the reduced invasiveness of ARL15-depleted primary RASFs aligns with the attenuated synovial pathology observed in vivo, collectively identifying ARL15 as a multifaceted regulator of synovial fibroblast pathogenicity.

*ARL15* expression partially overlapped with inflammatory fibroblast regions in RA synovium. Comparison of feature plots for *ARL15, IL6*, and *TNF* revealed enrichment of *ARL15* within inflammatory fibroblast clusters, particularly in the synovial sublining compartment (Figure 5). The spatial association between ARL15 and inflammatory marker expression supports a role for ARL15 in activated fibroblast states associated with RA pathogenesis.

In summary, our findings identify ARL15 as a potential therapeutic target in RA, with functional roles spanning inflammatory activation, fibroblast invasiveness, and arthritis severity. Future studies dissecting the molecular mechanisms linking ARL15 to inflammatory signaling, interferon responses, and cytoskeletal or trafficking pathways may further clarify how this small GTPase contributes to synovial fibroblast dysfunction and disease progression.

## References

1. Crofford LJ. Use of NSAIDs in treating patients with arthritis. Arthritis Research & Therapy. 2013;15(S3).

2. Albrecht K, Müller-Ladner U. Side effects and management of side effects of methotrexate in rheumatoid arthritis. Clin Exp Rheumatol. 2010;28(5 Suppl 61):S95–101.

3. Smolen JS, Aletaha D, Barton A, Burmester GR, Emery P, Firestein GS, et al. Rheumatoid arthritis. Nature Reviews Disease Primers. 2018;4(1):18001.

4. Bossaller L, Rothe A. Monoclonal antibody treatments for rheumatoid arthritis. Expert Opinion on Biological Therapy. 2013;13(9):1257–72.

5. Carmona L, Hernández-García C, Vadillo C, Pato E, Balsa A, González-Alvaro I, et al. Increased risk of tuberculosis in patients with rheumatoid arthritis. J Rheumatol. 2003;30(7):1436–9.

6. Okada Y, Wu D, Trynka G, Raj T, Terao C, Ikari K, et al. Genetics of rheumatoid arthritis contributes to biology and drug discovery. Nature. 2014;506(7488):376–81.

7. Negi S, Juyal G, Senapati S, Prasad P, Gupta A, Singh S, et al. A genome-wide association study reveals ARL15, a novel non-HLA susceptibility gene for rheumatoid arthritis in North Indians. Arthritis Rheum. 2013;65(12):3026–35.

8. Gillingham AK, Munro S. The small G proteins of the Arf family and their regulators. Annu Rev Cell Dev Biol. 2007;23:579–611.

9. Rocha N, Payne F, Huang-Doran I, Sleigh A, Fawcett K, Adams C, et al. The metabolic syndrome-associated small G protein ARL15 plays a role in adipocyte differentiation and adiponectin secretion. Scientific Reports. 2017;7(1).

10. Kapoor M, Wang J-C, Wetherill L, Le N, Bertelsen S, Hinrichs AL, et al. Genome-wide survival analysis of age at onset of alcohol dependence in extended high-risk COGA families. Drug and Alcohol Dependence. 2014;142:56–62.

11. Zegkos T, Kitas G, Dimitroulas T. Cardiovascular risk in rheumatoid arthritis: assessment, management and next steps. Therapeutic Advances in Musculoskeletal Disease. 2016;8(3):86–101.

12. Zhao J, Wang M, Deng W, Zhong D, Jiang Y, Liao Y, et al. ADP-ribosylation factor-like GTPase 15 enhances insulin-induced AKT phosphorylation in the IR/IRS1/AKT pathway by interacting with ASAP2 and regulating PDPK1 activity. Biochem Biophys Res Commun. 2017;486(4):865–71.

13. Kashyap S, Kumar U, Pandey AK, Kanjilal M, Chattopadhyay P, Yadav C, et al. Functional characterisation of ADP ribosylation factor-like protein 15 in rheumatoid arthritis synovial fibroblasts. Clin Exp Rheumatol. 2018;36(4):581–8.

14. Kashyap S, Pandey AK, Kumar P, Kanjilal M, Kumar U, Thelma BK. Dissecting ARL15 Function in Rheumatoid Arthritis: Insights From Ex Vivo and in Vitro Synovial Fibroblast Models. International Journal of Rheumatic Diseases. 2026;29(5):e70668.

15. Brand DD, Latham KA, Rosloniec EF. Collagen-induced arthritis. Nature Protocols. 2007;2(5):1269–75.

16. Trentham DE, Townes AS, Kang AH. Autoimmunity to type II collagen an experimental model of arthritis. J Exp Med. 1977;146(3):857–68.

17. Hafner M, Schmitz A, Grüne I, Srivatsan SG, Paul B, Kolanus W, et al. Inhibition of cytohesins by SecinH3 leads to hepatic insulin resistance. Nature. 2006;444(7121):941–4.

18. Zhu W, London NR, Gibson CC, Davis CT, Tong Z, Sorensen LK, et al. Interleukin receptor activates a MYD88-ARNO-ARF6 cascade to disrupt vascular stability. Nature. 2012;492(7428):252–5.

19. Smith MH, Gao VR, Periyakoil PK, Kochen A, Dicarlo EF, Goodman SM, et al. Drivers of heterogeneity in synovial fibroblasts in rheumatoid arthritis. Nature Immunology. 2023;24(7):1200–10.

20. Kashyap S, Chattopadhyaay P, Pandey AK, Kumar U, Yadav CS, Thelma B, editors. ARL15 expressed by rheumatoid arthritis synovial fibroblasts regulates IL6 and plays a role in apoptosis and hypoxia. NARTHRITIS & RHEUMATOLOGY; 2016: WILEY 111 RIVER ST, HOBOKEN 07030–5774, NJ USA.

21. Guo H, Callaway JB, Ting JPY. Inflammasomes: mechanism of action, role in disease, and therapeutics. Nature Medicine. 2015;21(7):677–87.

22. Firestein GS, McInnes IB. Immunopathogenesis of Rheumatoid Arthritis. Immunity. 2017;46(2):183–96.

23. Lubberts E. The IL-23–IL-17 axis in inflammatory arthritis. Nature Reviews Rheumatology. 2015;11(7):415–29.

24. McInnes IB, Buckley CD, Isaacs JD. Cytokines in rheumatoid arthritis — shaping the immunological landscape. Nature Reviews Rheumatology. 2016;12(1):63–8.

25. Notley CA, Inglis JJ, Alzabin S, McCann FE, McNamee KE, Williams RO. Blockade of tumor necrosis factor in collagen-induced arthritis reveals a novel immunoregulatory pathway for Th1 and Th17 cells. The Journal of Experimental Medicine. 2008;205(11):2491–7.

