## Supplementary material for "*ARL15* promotes inflammatory fibroblast activation and disease severity in rheumatoid arthritis: integrated transcriptomic and collagen-induced arthritis model analyses"

**Bacterial cloning**

The commercially procured *ARL15* plasmid (RC205610, Origene, USA) was resuspended in molecular biology grade water and quantified using a NanoDrop spectrophotometer. Transformation was performed using chemically competent E. coli Dh5α cells, followed by selection of transformants on LB agar plates containing 25 μg/ml Kanamycin. Overnight cultures of positive clones were grown at 37°C with shaking at 250 rpm. Small aliquots of all cultures were stored at 4°C, while the remaining overnight cultures (~2.5 ml) were used for plasmid DNA isolation using a miniprep protocol involving cell lysis with cell lysis buffer, neutralization solution, and purification through PureYield™ Minicolumns (Promega, USA), including washing steps to remove endotoxins and contaminants. DNA was eluted in 30 μl elution buffer, and quality and quantity were confirmed by NanoDrop. All clones were screened by single and double restriction enzyme digestions using BamH1 and Xho1 enzymes, incubated overnight at 37°C, and analyzed on 2% agarose gels alongside uncut plasmids and DNA ladders. Sanger sequencing was performed on plasmids from all clones using the *ARL15* forward primer (5’CATCAAGACAA GCCAGCAGC3’), with DNA samples diluted to 500 ng in 5 μl volume. The positive clone cultures harboring the correct plasmid were expanded, and plasmid preparations were performed in larger volumes using the same protocol. The pooled DNA was stored at −20°C for subsequent use in MH7A cell transfection.

**MH7A Transfection**

Transfection of MH7A cells (3 different passages) was conducted using a standard liposome-mediated approach with Lipofectamine 2000 (11668019, ThermoFisher Scientific, USA). Cells were plated to reach 70-90% confluence on the day of transfection. One hour prior to transfection, complete medium was replaced with Opti-MEM I reduced serum medium (31985070, ThermoFisher Scientific, USA). Transfection complexes were prepared by diluting 500 ng of ARL15 plasmid DNA and Lipofectamine 2000 separately in Opti-MEM I medium, combined, and incubated for 15 minutes at room temperature. Complexes were added to MH7A cells and incubated at 37°C with 5% CO2 for 6 hours. After this incubation, the Opti-MEM medium was replaced with complete medium, and cells were further incubated at 37°C with 5% CO2 for 48 hours before RNA isolation. A Lipofectamine-only control served as the negative experimental control.

RNA isolation was performed using the Direct-zol RNA isolation kit (R2070, Zymo Research, USA), which allows direct application of samples in TRI Reagent® to the spin column for RNA purification without phase separation. Extracted RNA was quantified using a fluorometry-based RNA quantification kit (Q32852, Thermo Fisher Scientific, USA). RNA samples thus isolated were used for Real-Time PCR experiments (using Actin UBC as reference controls) and transcriptome sequencing. The overexpression confirmed samples in triplicate (3 overexpression and 3 lipofectamine control samples) were used for transcriptome sequencing.

**Transcriptome sequencing**

Transcriptome sequencing for all samples was conducted at a commercial facility (MedGenome, Bangalore, India). At the commercial facility, all samples were quantified using the fluorometry-based RNA quantification kit (Q10211, ThermoFisher Scientific, USA), RNA purity was assessed with QIAxpert (Qiagen, USA), and RNA integrity was evaluated on a TapeStation using RNA screenTapes (5067-5576, Agilent, USA). Libraries were prepared using the NEB Ultra II directional RNA-Seq Library Prep Kit protocol (E7760L, NEB, USA)	. Ribosomal RNAs (~95% of total RNA) were removed from 100 ng total RNA by hybridization with biotinylated, target-specific oligos and Ribo-Cop rRNA removal beads. Ribodepleted RNA was fragmented with divalent cations at elevated temperature, reverse transcribed to first strand cDNA, then second strand synthesis was performed using DNA Polymerase I and RNase H. The cDNA was end-repaired, 3’ A-tailed, and adapters were ligated. Adapter-ligated products were purified and enriched by PCR (initial denaturation 98°C 30s; 13 cycles of 98°C 10s, 65°C 75s; final extension 65°C 5 min). PCR products were purified and checked for size distribution on Fragment Analyzer with HS NGS Fragment Kit (DNF-474-1000, Agilent, USA). Libraries were quantified with fluorometry-based RNA quantification kit (Q32852, Thermo Fisher Scientific, USA), pooled, and sequenced on Illumina NovaSeq 6000 to generate 150 bp paired-end reads.

**RNA isolation**

RNA isolation was performed using the Direct-zol RNA isolation kit (R2070, Zymo Research, USA) and quantified using a fluorometry-based RNA quantification kit (Q32852, Thermo Fisher Scientific, USA). RNA samples thus isolated were used for Real-Time PCR experiments using Actin UBC as reference controls. Overexpression was confirmed by qPCR and samples in triplicate (overexpression and lipofectamine controls) were used for transcriptome sequencing.

**Bioinformatic analysis**

Bioinformatic analysis was performed for all samples, beginning with a quality assessment of the raw FASTQ sequencing data. This included evaluation of base quality score distributions, overall sequence quality, average base content, GC percentage, rates of PCR duplicates, over-represented sequences, and potential adapter contamination. Subsequently, Trimmomatic (v0.36, <https://doi.org/10.1093/bioinformatics/btu170>) was used to trim adapter sequences and remove low-quality bases, ensuring that only high-quality reads were retained for all downstream analysis. Contamination removal was accomplished with Bowtie2 (v2.2.4, PMID: [22388286](https://pubmed.ncbi.nlm.nih.gov/22388286/)), effectively filtering out non-polyA RNA species such as mitochondrial genome sequences, rRNAs, tRNAs, and any residual adapters. The curated paired-end reads were then aligned to the human reference genome (hg19) using HISAT2 (v2.1.0, <https://www.nature.com/articles/nmeth.3317>) to generate accurate mapping data for quantification. Gene expression quantification was conducted using FeatureCount (v1.5.2, <https://doi.org/10.1093/bioinformatics/btt656>), resulting in raw read count matrices that were normalized using DESeq2 (DOI: [10.1186/s13059-014-0550-8](https://doi.org/10.1186/s13059-014-0550-8)) to correct for library size and composition. Differential expression analysis was carried out via DESeq2 normalization, calculation of fold changes between treated and control samples, and selection of genes based on a significance threshold of p ≤ 0.05. Genes exhibiting a log2(fold change) of ≤ −1 or ≥ 1 were classified as statistically significant and prioritized for further biological interpretation and pathway analysis.

**CIA induction**

DBA1/J male mice obtained for the study were randomly divided into 9 experimental groups and marked by animal accession number, cage card and permanent tail mark. On day 1 after acclimatization of 5 days, all DBA/1 mice except those in the healthy group were immunized by intradermal injections of an emulsion containing 100 μg of immunization grade bovine type II collagen in complete Freund's adjuvant containing heat-killed Mycobacterium butyricum the base of the tail. On day 21, a booster containing 100 μg bovine CII in incomplete Freund's adjuvant was administered following previously described protocol (15). The animals were being observed for disease development. Drugs were administered as described in Supplementary table 1. The healthy group did not receive any treatment. A schematic presentation of the disease induction and treatment protocol is shown in Figure 1A. All the groups of mice were regularly scored for disease indicators (inflammation, redness, severity etc) according to their severity for 2 weeks until termination. Bone and tissue sections samples were collected from the experiment animals as detailed earlier. All the reagents used for the generation of the CIA models were obtained from M/s Chondrex Inc., USA.

**Clinical data collection**

Clinical score data were recorded during the study period. Mean clinical score change, percentage change between each treatment group at different time points or at the end of the study period were calculated and compared with respective vehicle treatment (Supplementary table 5).

**Collection of bones and tissues from experimental animals**

Prior to killing of the mice at the end of the experimental schedule, 14 days after initiation of respective treatment regimen (Figure 1A), 100-300 µl of blood/ mouse was drawn from the retino-orbital sinus of all the mice and used for biomarker analyses. Long bones including the joints were denuded and stored in formalin.

**Measurement of serum cytokines**

The sera samples were diluted as per manufacturer’s instruction and 100µl of samples along with the diluted standards were loaded in the well of ELISA strips. Biotin conjugate was then added to each of the wells followed by incubation for 2 hours at room temperature. Color was developed in the wells by treatment of streptavidin-HRP followed by TMB substrate solution. OD readings in each of the wells were captured at 450nm on spectrophotometer (Tecan, Switzerland) and the concentrations of cytokines were derived based on the values of standards. A five-point curve fit was drawn using the optical density (OD) values obtained for standards. The OD value for the unknown samples was then extrapolated using Elisa online tool to find the final concentration of cytokines.

**Results**

**Cytokine and Antibody Levels**

**IgG1 Antibody**: The mean serum IgG1 level in the Vehicle group was 128367.88. The groups treated with AntiARL15 antibody mAb 1 (86433.87 ng/mL), AntiARL15 antibody BSA and Azide free (64246.86 ng/mL), and the combination (96369.68 ng/mL) showed visual reductions in IgG1 levels compared to the Vehicle and Isotype controls (G3, mean 146,290.48 ng/mL) (Supplementary Figure 2B,C).

**IL-6 and IL-1β**: Pro-inflammatory cytokine levels were generally high in the disease control groups. The mean IL-6 level for the Vehicle group was 2,434.73 pg/mL, whereas for the isotype it was 3449.71 pg/mL. Treatment groups generally had much lower mean IL-6 levels AntiARL15 mAb 1 at 107.82 pg/mL, AntiARL15 mAb 2 at 73.10 pg/mL,the combination at 25.72 pg/mL and for the Isoquinoline group it was 198.26 pg/mL. We did not find any significant difference in the siRNA group. Similarly, IL1-β levels were found to be high in the vehicle group at 127.19 pg/mL and isotype group at 138.31 pg/mL. The combination group (78.39 pg/mL) of antibodies showed the maximum decrease in IL1-β level upon treatment, whereas the mAb1 and mAb2 remained at 106.87 pg/mL and 106.07 pg/mL. IL1β decreased slightly in the Isoquinoline group (111.70 pg/mL) whereas there was no significant change in IL1β levels between the control and disease siRNA group (Supplementary table7).

**TNFα and IL17**: The levels of TNFα remained unchanged in the siRNA control and ARL15 siRNA group but there was a significant reduction in TNFα levels in the G4 (23.27 pg/mL), G5 (16.45 pg/mL), G6 (6.16 pg/mL) and G9 (26.94 pg/mL) at as compared to the disease groups G2 (480.16 pg/mL) and G3 (1297.79 pg/mL). IL17 levels were not significant in either of the disease or treatment groups (Supplementary table7).
