## Supplementary figures and images for "*ARL15* promotes inflammatory fibroblast activation and disease severity in rheumatoid arthritis: integrated transcriptomic and collagen-induced arthritis model analyses"

### Supplementary Figure 1

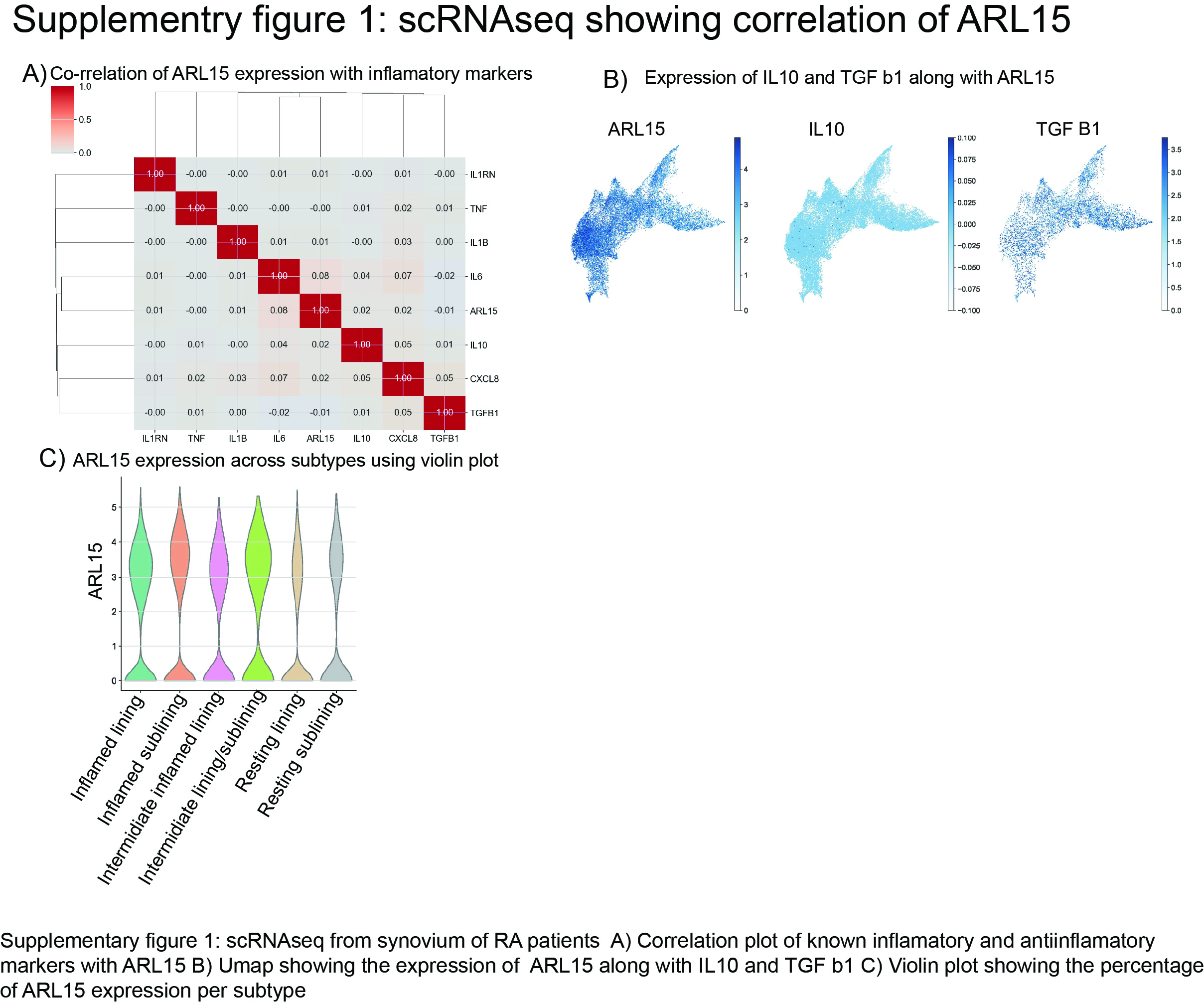

### Supplementary Figure 2

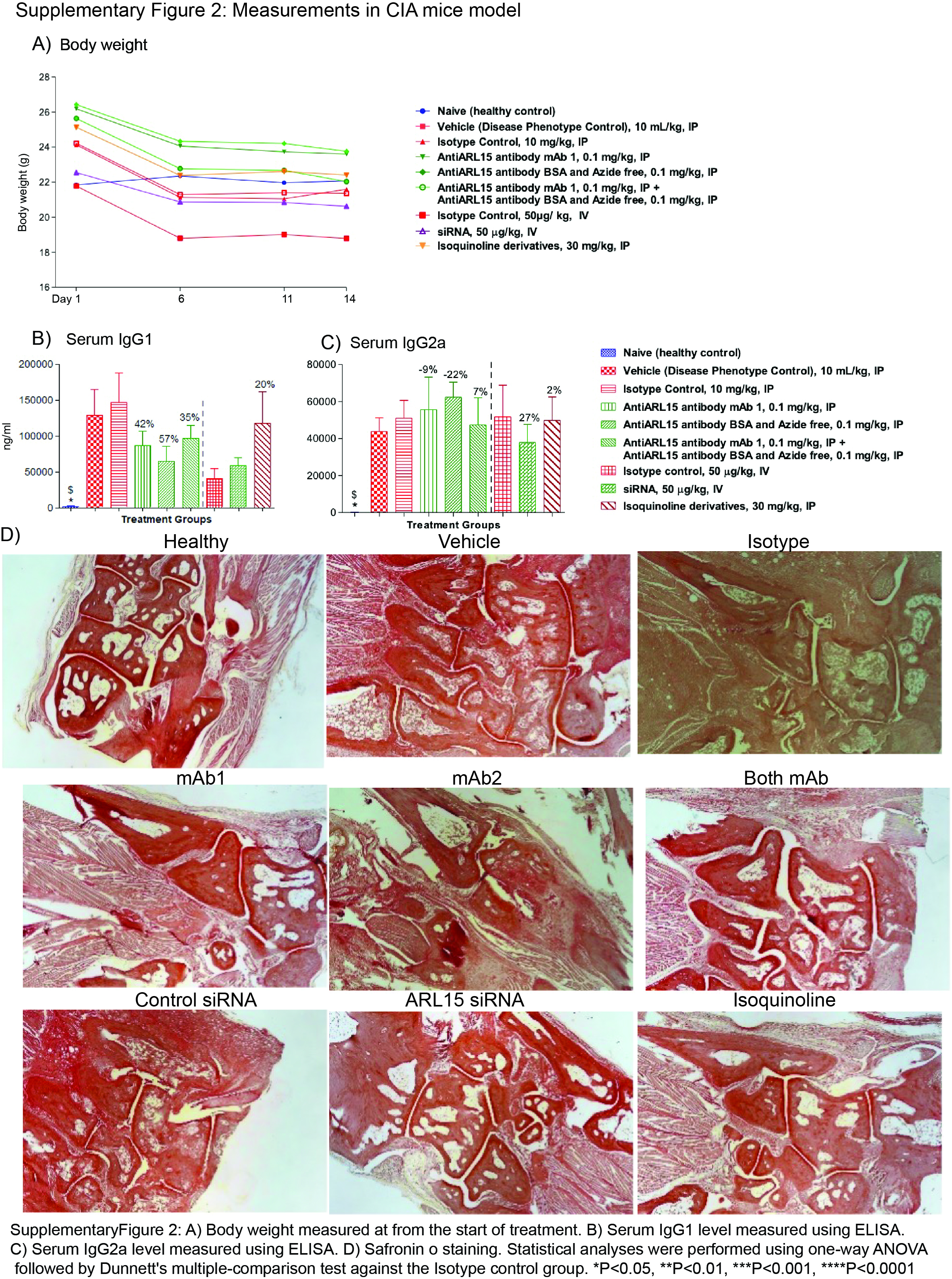
